# Notch receptors involved in the choice between intestinal secretory and enterocytes and differentiation of Bestrophin 4 cells

**DOI:** 10.64898/2025.12.17.694952

**Authors:** Samah Allayati, Pijush Sutradhar, Morgan Prochaska, Lea Maney, Christian Choy, Abrielle Swartz, Kenneth Wallace

## Abstract

During the first half of embryogenesis, the intestinal epithelium is a simple layer of cuboidal cells surrounded by lateral plate splanchnic mesoderm. As embryogenesis proceeds, the epithelium differentiates to include a variety of enterocytes and secretory cells. During the period of epithelial cell differentiation a number of signaling pathways are used to generate the full complement of cell types. Notch signaling is important for the choice between enterocytes and secretory cells. Following the initial choice, both enterocytes and secretory cells continue differentiating into a number of different subtypes of each cell type. In the zebrafish embryo, Notch signaling is utilized at least three times during development of the intestinal epithelium. The first two Notch signaling events are involved in the choice between enterocytes and secretory cells, while the third involves differentiation of a unique cell type that we previously called Notch Receiving Secretory Cells (NRSCs). While we have identified the times when Notch signaling is active, the individual Notch receptors used in each signaling event have not been previously identified. Here we use loss of function mutants in the four Notch receptors (*notch 1a, 1b, 2*, and *3*) to identify which are used in the each of the Notch signaling events. We find redundant use of Notch receptors in the choice between enterocytes and secretory cells with loss of combinations of either *notch 2* and *notch 3* or *notch 2* and *notch 1b* but not loss of individual receptors resulting in changes to secretory cell numbers. During the Notch signaling event following the choice between secretory and enterocytes, we identify NRSCs as Bestrophin 4 (BEST4+) cells. In contrast to the choice between secretory and enterocyte cells, we find that loss of individual Notch receptors result in differing numbers of BEST4+ cells. Even though loss of individual receptors affect BEST4+ numbers we suggest that combinations of Notch receptors play a role in differentiation of different BEST4+ subtypes.

**Highlights:**

- Requirements for the choice between secretory and enterocytes within the intestinal epithelium require Notch2/Notch3 and Notch2/Notch1b combinations.
- Notch 2 and Notch1b are required for Bestropin 4 (BEST4+) cell development.
- Combinations of Notch2, Notch1b, and Notch3 may play roles in formation of different BEST4+ subtypes.

## Introduction

By the end of embryogenesis, the zebrafish intestine differentiates into an organ which is able to ingest and process food to provide energy for the organism (Westerfield, 1993). Early in embryogenesis, the vertebrate intestinal epithelium is a simple layer of cuboidal cells surrounded by lateral plate splanchnic mesoderm (Roberts, 2000). As the intestine develops, epithelial cells become columnar as they differentiate. Surface area of the epithelium increases by either forming broad folds as in zebrafish(Wallace et al., 2005) or individual villi in mammalian species(Noah et al., 2011). Each intestine develops a proliferative stem cell region at the base of the folded structure(Noah et al., 2011; Wallace et al., 2005). In the adult intestine, progeny from the stem cell region move towards the tip, undergo apoptosis, and are extruded into the lumen(Noah et al., 2011; Wallace et al., 2005).

Halfway through zebrafish embryogenesis, intestinal epithelial cells are specified as either enterocytes or secretory cells, utilizing Notch lateral inhibition (Flasse et al., 2013; Roach et al., 2013). Lateral inhibition utilizes Notch signaling to regulate the numbers of two different cell types within an epithelium (Zhou et al., 2022). In the zebrafish intestinal epithelium, cells that take on the secretory fate, upregulate *ascl1a* and the DeltaD ligand. Delta D ligands bind to Notch receptors, inducing the cell with the activated Notch receptor to enter an enterocyte fate (Flasse et al., 2013; Roach et al., 2013). If *ascl1a* is absent, no epithelial cells enter the secretory cell fate (Flasse et al., 2013; Roach et al., 2013). When *ascl1a* is present but there is an absence of Notch signaling, the number of epithelial cells entering the secretory pathway increases.

Within the zebrafish intestine, we find at least three different periods of where Notch signaling is active (Roach et al., 2013). The first two periods involve the choice between secretory and enterocyte cells. The first period occurs between 30 to 34 hpf, but is unclear as to how Notch signaling affects numbers of secretory cells as this appears to be before the lateral inhibition phase. The second period of Notch signaling occurs between 64 to 74 hpf. This second phase occurs when there is active choice between secretory and enterocytes, coinciding with expression of other components needed for lateral inhibition; *ascl1a* and *deltaD*. Inhibition of Notch signaling, using the gamma secretase inhibitor DAPT, during either 30 to 34 or 64 to 74 hpf results in increased numbers of secretory cells (Roach et al., 2013). The third period of Notch signaling does not appear to involve lateral inhibition but instead involves formation of a unique secretory cell subtype which begins differentiating after 74 hpf (previously referred to as Notch Receiving Secretory Cells-NRSCs). New NRSCs continue to differentiate throughout the post embryonic period(Li et al., 2019). We previously visualized NRSCs using a Notch driven Cre ERT2 combined with a nuclear mCherry reporter (referred to as Ncre) (Li et al., 2019; Wang et al., 2011). Inhibition of Notch signaling during the period when most NRSCs form (between 74 to 76 hpf) does not alter the overall number of secretory cells, suggesting the choice between enterocytes and secretory cells is completed (Li et al., 2019). As we demonstrate here, Notch signaling during this period plays a role in differentiation of NRSCs.

While Notch signaling is active during at least three periods of intestinal development, we do not know which of the Notch receptors participate in these events. Zebrafish have Notch 1a/1b, Notch 2, and Notch 3 while mammals have Notch 1, 2, 3, and 4 receptors. (Bierkamp and Campos-Ortega, 1993; Westin and Lardelli, 1997; Zhou et al., 2022) In the mammalian intestine, Notch 1 and 2 are expressed within the adult small intestinal epithelial crypt cells while Notch 3 and 4 are expressed in surrounding endothelial and mesenchymal cells (Fre et al., 2011; Riccio et al., 2008). The mammalian Notch 1 and Notch 2 appear to act redundantly, as inactivation of both receptors results in conversion of the crypt to post mitotic goblet cells while inactivation of only Notch 1 or Notch 2 produces no phenotype (Fre et al., 2011).

In contrast to mammals, all four zebrafish Notch receptors are expressed throughout the intestine (Lorent et al., 2004), suggesting that any of the receptors may be active during identified signaling periods. To determine which of the Notch receptors are involved in each signaling event we use two assays. Loss of all Notch signaling using DAPT results in increases in secretory cell numbers. In the first assay, loss of function of the correct combination of Notch receptors will also produce increased secretory cell numbers as with inhibition of all Notch receptors using DAPT. In the second assay to identify receptors involved in formation of the NRSCs, loss of individual or combinations of Notch receptors should result in low numbers or loss of cells expressing the Notch reporter (combination of CreERT2 driven by activated Notch receptor expressing mCherry fluorescence-Ncre) throughout the epithelium. Due to differences in use of Notch receptors between these two events, we have been able to sort out different use of Notch receptors involved in secretory/enterocyte cell choice and differentiation of NRSCs.

As we demonstrate in this work, the choice between secretory and enterocyte cells relies on combinations of Notch receptors. Individual mutants do not result in statistically significant changes in the number of secretory cells. The number of NRSCs in contrast rely more on individual Notch receptors. Even though loss of individual Notch receptors reduce NRSC numbers, we find that multiple Notch receptors may be required for differentiation of subtypes of this cell type. Recent single cell sequencing identifies Notch signaling within a unique epithelial cell type of Bestrophin 4 (BEST4+) cells which are defined by expression of genes such as Notch2, BEST4, SPIB, CA7, and CFTR (Burclaff et al., 2022; Busslinger et al., 2021; Elmentaite et al., 2021; Hickey et al., 2023; Parikh et al., 2019; Smillie et al., 2019; Sur et al., 2023a; Wang et al., 2025; Willms et al., 2022). We find that the cells we call NRSCs are BEST4+ cells. Previously, we interrupted Notch signaling in NRSCs/ BEST4+ cells using DAPT (inhibiting cleavage of the Notch Intracellular Domain-NICD) and found increased epithelial proliferation (Li et al., 2019). *notch2* mutants with the loss of all BEST4+ cells, also have increased epithelial proliferation.

## Results

### Notch receptors 2, 3, and 1b act redundantly during the choice between secretory and enterocyte cell formation

Most epithelial cells within the intestine differentiate into either an enterocyte lineage or a secretory lineage(Kolev and Kaestner, 2023). Following this initial decision, enterocytes and secretory cells continue differentiating into numerous subtypes (Capdevila et al., 2021). As with other intestines, the zebrafish epithelium utilizes Notch signaling to regulate the number of cells entering the secretory fate (Crosnier et al., 2005). Cells that will ultimately take on a secretory fate, increase expression of the *ascl1a* and the Notch ligand DeltaD (Roach et al., 2013). Surrounding cells activate Notch signaling to differentiate into enterocyte cells. Without Notch signaling, surrounding cells will develop into secretory cells and not enter the enterocyte fate. As a result, reduction of Notch activation during times when the signaling is active, results in increased secretory cell numbers (Roach et al., 2013). With reduction in Notch activity using the gamma secretase inhibitor, (2*S*)-*N*-[(3,5-Difluorophenyl)acetyl]-L-alanyl-2-phenyl]glycine 1,1-dimethylethyl ester (DAPT), we have shown that the initial decision to differentiate into epithelial or secretory fates occurs during two periods during embryogenesis. The first is between 30-34 hpf and the second is between 64 to 74 hpf (Roach et al., 2013).

While Notch signaling is active at multiple times during embryonic intestinal development, which receptors are activated during each of these signaling events are not known. To determine which receptors are utilized in the choice between secretory and enterocytes, we use Notch loss of function mutants for each of the four receptors (Notch1a, 1b, 2, and 3). As with Notch inhibition following DAPT exposure (Roach et al., 2013), loss of the Notch receptor or combination of receptors responsible for signaling should increase secretory cell numbers. Using null mutants for each of the four Notch receptors, we performed dihybrid crosses for each combination of mutants. For each combination of Notch receptor mutants we genotyped embryos from each cross to obtain double mutants, single mutants, and WT embryos. A pan secretory cell antibody to annexin A4 (2F11) (Crosnier et al., 2005; Roach et al., 2013; Zhang et al., 2014) was used to sample secretory cell numbers in both an anterior and posterior region of each of the embryonic intestines (sampling regions shown in Figure 1A).

**Figure 1.**
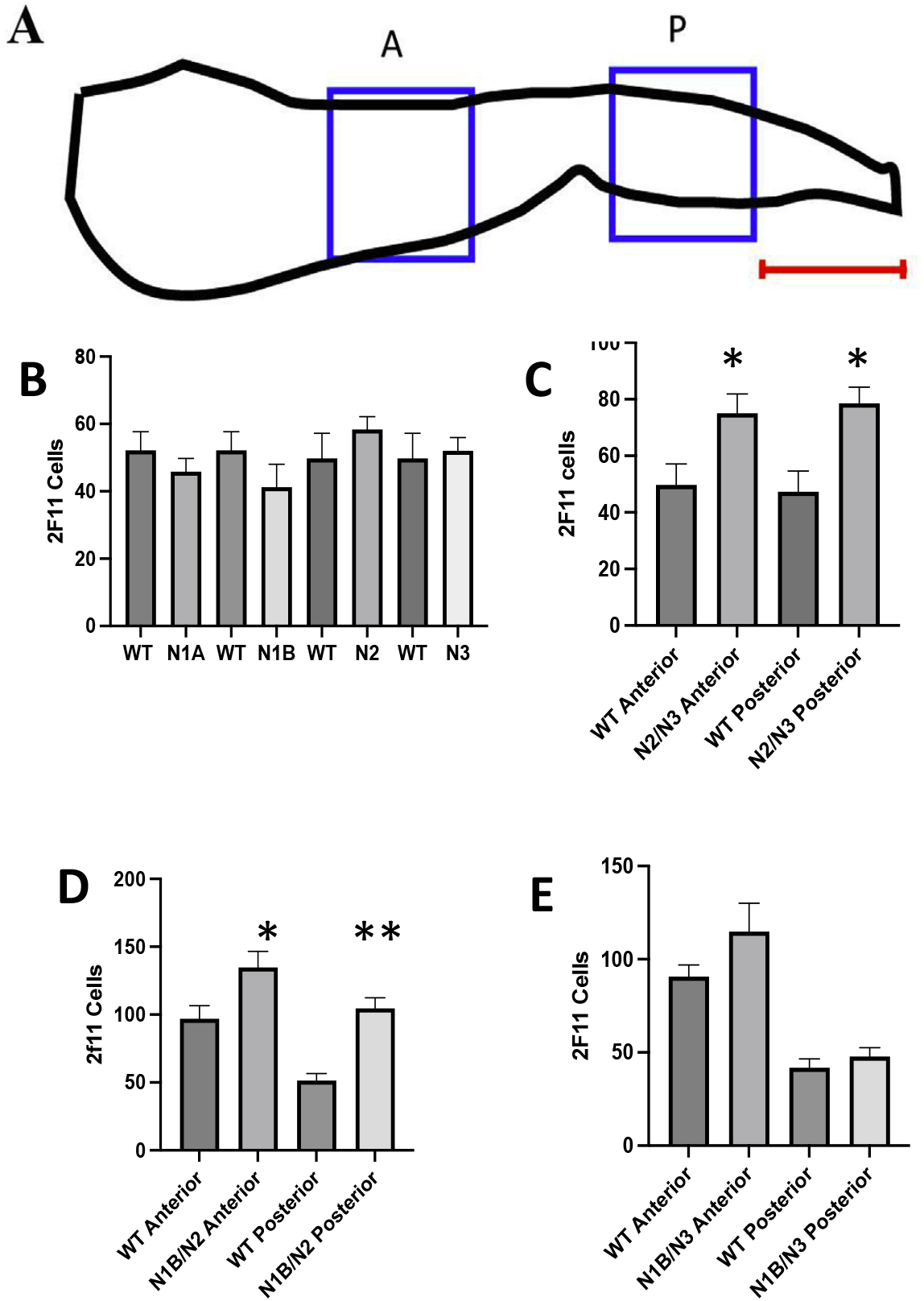
Changes in secretory cell numbers due to Notch receptor mutants. A. Diagram of regions recorded in anterior (A blue box) and posterior (P blue box) intestine. Each region is 183 X 183mm. The anterior region is where the intestinal bulb begins to narrow. The posterior region is 183 mm from the posterior end (indicated by red line). Mutants and WT siblings are paired in each panel. B. Number of AnnexinA4 (2F11) expressing cells in anterior region of WT siblings compared to mutant Notch receptors. C. Number of AnnexinA4 (2F11) expressing cells in *notch2/notch3* double mutants compared to WT siblings. D. Number of AnnexinA4 (2F11) expressing cells in *notch1b/notch2* double mutants compared to WT siblings. E. *notch1b/notch3* double mutants compared to WT siblings. Single notch mutants n=5 to 8; notch 2/notch3 n= 7; *notch1b*/*notch2* mutants n= 8 to 10; *notch 1b*/*notch3* n=6 to 10; * p value < 0.05; ** p value < 0.01; *** p value < 0.001

We first compared the number of secretory cells in embryos with individual mutant Notch receptors to WT at the end of embryogenesis (5 dpf). None of the single Notch receptor mutants show any statistically significant alteration in secretory cells (Figure 1B). We next compared changes in the number of secretory cells in double mutant combinations. We are able to obtain double mutants from all of the dihybrid crosses, except from combinations that include *notch 1a*. As *notch 1a* null mutants also produce additional neural and somite defects (Holley et al., 2002), loss of *notch1a* combined with additional Notch receptors appear to cause lethality. Out of the remaining group of double mutant combinations, N2 and N3 (Figure 1C) or N2 and N1B (Figure 1D) are the pairs that demonstrate a statistically significant increase in secretory cell numbers. Loss of the combination of N1B and N3 does not result in statistically significant changes in secretory cell numbers (Figure 1E). Increased secretory cell numbers only with double Notch receptor mutant combinations suggest that Notch receptors work redundantly in the choice between secretory and enterocytes. Single Notch receptor mutants do not result in changes to secretory and enterocyte cell numbers.

### Notch receptors are activated in BEST4+ cells beginning on the third day of embryogenesis

We previously identified a subtype of secretory cells that receive Notch signaling (Notch receiving secretory cells-NRSCs) (Li et al., 2019). These cells begin differentiating at 74 hpf, following the choice between enterocytes and secretory cells using a new round of Notch signaling (Li et al., 2019). New NRSCs continue differentiating through the end of embryogenesis and throughout the post embryonic period (Li et al., 2019). We had not further identified these cells beyond the fact that they are part of the secretory lineage and have activated Notch receptors.

BEST4+ cells are a recently identified unique intestinal epithelial cell type, first identified in humans, that expresses genes such as Notch2, BEST4, SPIB, CA7, and CFTR (Burclaff et al., 2022; Busslinger et al., 2021; Elmentaite et al., 2021; Hickey et al., 2023; Parikh et al., 2019; Smillie et al., 2019; Sur et al., 2023a; Wang et al., 2025; Willms et al., 2022). BEST4+ cells are present in the zebrafish, rat, pig, and rabbit intestinal epithelium model systems but absent in mouse (Malonga et al., 2024). Due to BEST4+ cells expressing *notch2* receptor and identification of NRSCs through activated Notch, we hypothesized that NRSCs are BEST4+ cells.

To determine whether NRSCs are BEST4+ cells, we co-localized cells with activated Notch receptors and *best4* expression. As previously demonstrated, *best4* is expressed in single cells from the anterior to posterior intestine (Sur et al., 2023b; Willms et al., 2022). To identify cells with activated Notch receptors, we used the transgene *Tg(T2KTp1glob:creER*^*T2*^ *)jh12* which drives CreER^T2^ using *12 RBP-Jκ-binding sites* and a minimal β-globin promoter. The Cre ER^T2^ driver is combined with the transgene *Tg(T2Kβactin:loxP-stop-loxP-hmgb1-mCherry)*^*jh15*^ to express nuclear mCherry in cells with activated Notch receptors (Wang et al., 2011). We refer to the combination of the two transgenes as Ncre. Embryos containing Ncre were induced with 4OHT at 74 hpf and grown to 5 dpf. We find that *best4* (Figure 2A) expression colocalizes with nuclear mCherry (Figure 2B and 2C). We find that about 40% of the *best4* cells express nuclear mCherry, but we do not find any nuclear mCherry cells without *best4*.

**Figure 2.**
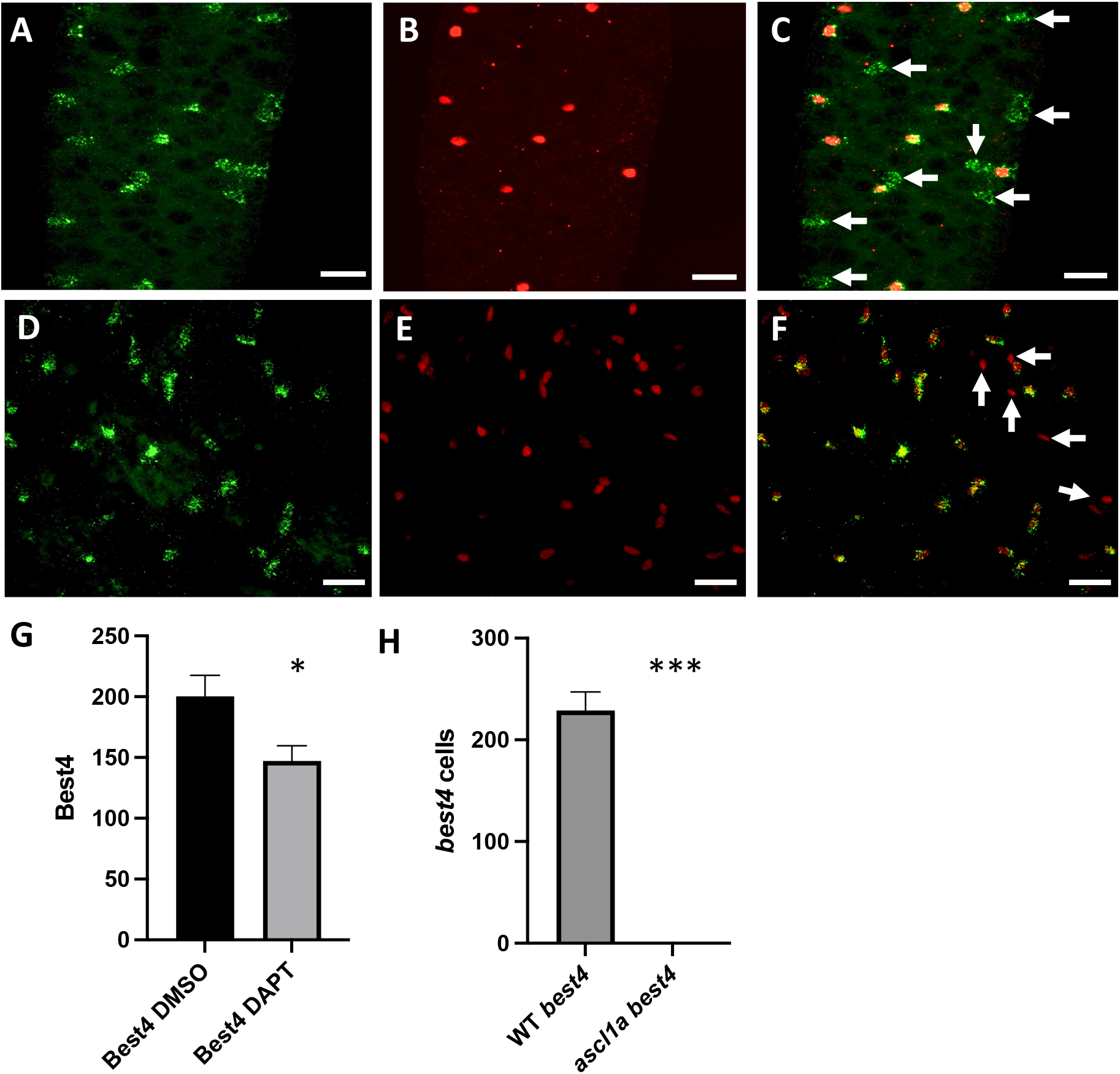
Identification of NRSCs as BEST4+ cells. A. *best4* RNA *in situ* hybridization (green) B. Nuclear mCherry (red-Notch activated reporter). C. Co-localization of *best4* and nuclear mCherry. Arrows indicate *best4* cells without nuclear mCherry expression. D. *best4* RNA *in situ* hybridization (green) E. Pan-secretory cell marker AnnexinA4 (red-2F11 immunohistochemistry). F. Co-localization of *best4* and pan secretory cell marker AnnexinA4. AnnexinA4 is also expressed in cells of the intrapancreatic ducts of the pancreas. Some of the intrapancreatic duct cells of the pancreas express AnnexinA4 to the side of the intestine with no *best4* expression (arrows). G. Number of *best4* cells (entire intestine) comparing DMSO control to DAPT treated embryos. H. Number of *best4* cells (entire intestine) comparing WT to *ascl1a* mutants. DAPT n=9; *ascl1a* n=8; Scale bar: 20 μM. ^*^ p value ≤ 0.05; ^**^ p value ≤ 0.01; ^***^ p value ≤ 0.001

We previously used the Notch inhibitor, DAPT, to interrupt development of *best4* cells (Li et al., 2019). At the time, we did not know the cells were *best4* and did not have an additional marker to determine the effect of DAPT treatment on the cell type. With identification of these cells as *best4*, we use *best4* RNA *in situ* hybridization to determine whether DAPT alters the number of *best4* cells within the epithelium. We applied DAPT to embryos for a similar time as before, 74 to 76hpf and grew the embryos to 5 dpf. Five day embryos treated with DAPT during the third day demonstrate a statistically significant reduction in *best4* cells (Figure 2G). While the reduction is statistically significant (p= 0.025), many *best4* cells remain.

There has been question about the lineage of BEST4+ cells as to whether they arise from a secretory or enterocyte lineage (Malonga et al., 2024). We previously found that all NRSCs express the pan-secretory cell marker AnnexinA4 (2F11 antibody) (Li et al., 2019) suggesting BEST4+ cells have a secretory origin. Ncre nuclear mCherry expression however, only represents about 40% of the *best4* cells. To confirm that even the *best4* cells without mCherry expression also co-localize with AnnexinA4, we combined *best4* RNA *in situ* with AnnexinA4 immunohistochemistry. We find that all *best4* cells (Figure 2D) co-localize with AnnexinA4 (Figure 2E and 2F) suggesting that even *best4* cells without mCherry expression arise from a secretory origin.

To further support the secretory nature of BEST4+ cells, we investigated expression of *best4* in *ascl1a* mutants. We previously identified a total lack of secretory cells in *ascl1a* null mutants (Roach et al., 2013). If BEST4+ cells are derived from the secretory cell lineage, then *ascl1a* mutants will lack all BEST4+ cells. We find no *best4* expression in *ascl1*a mutants (Figure 2H) further supporting that BEST4+ cells are derived from a secretory cell lineage.

### Notch 2 and Notch1b play roles in differentiation of BEST4+ cells

Following co-localization of Notch signaling with *best4* expressing cells in the intestinal epithelium, we asked which of the Notch receptors are necessary for BEST4+ cell development. We used Notch receptor null mutants in combination with Ncre transgenes to identify what individual or combination of Notch receptors demonstrate losses of BEST4+ cells. Loss of Notch receptors that play a role in development of BEST4+ cells should result in loss of the nuclear mCherry reporter in the intestinal epithelium.

We performed dihybrid crosses to create loss of function combinations of the Notch receptors. Embryos were analyzed at 5 dpf for the presence or absence of Notch signaling within the intestinal epithelium (presence or absence of nuclear mCherry). For each combination, embryos were genotyped to identify double mutant, single mutant, and WT embryos.

In contrast to the redundancy of Notch receptors in the choice between enterocytes and secretory cells, we find that individual Notch receptor mutants significantly reduce the number of mCherry cells. While there is an average of 118 cells within the epithelium that receive Notch signaling, *notch 2* mutants reduce the number of nuclear mCherry cells to an average of 0.33, *notch 3* mutants have a reduction to an average of 11.7 cells (Figure 3A), and *notch1b* reduces cells to an average of 47.7 (Figure 3B). There are no statistically significant changes in mCherry cell numbers in *notch1a* mutants (Figure 3B). The *notch2* /*notch3* mutant combination has a few more cells on average than the individual mutations (Figure 3A). In the *notch2*/*notch1b* combination, we find no nuclear mCherry cells within the epithelium (Figure 3C).

**Figure 3.**
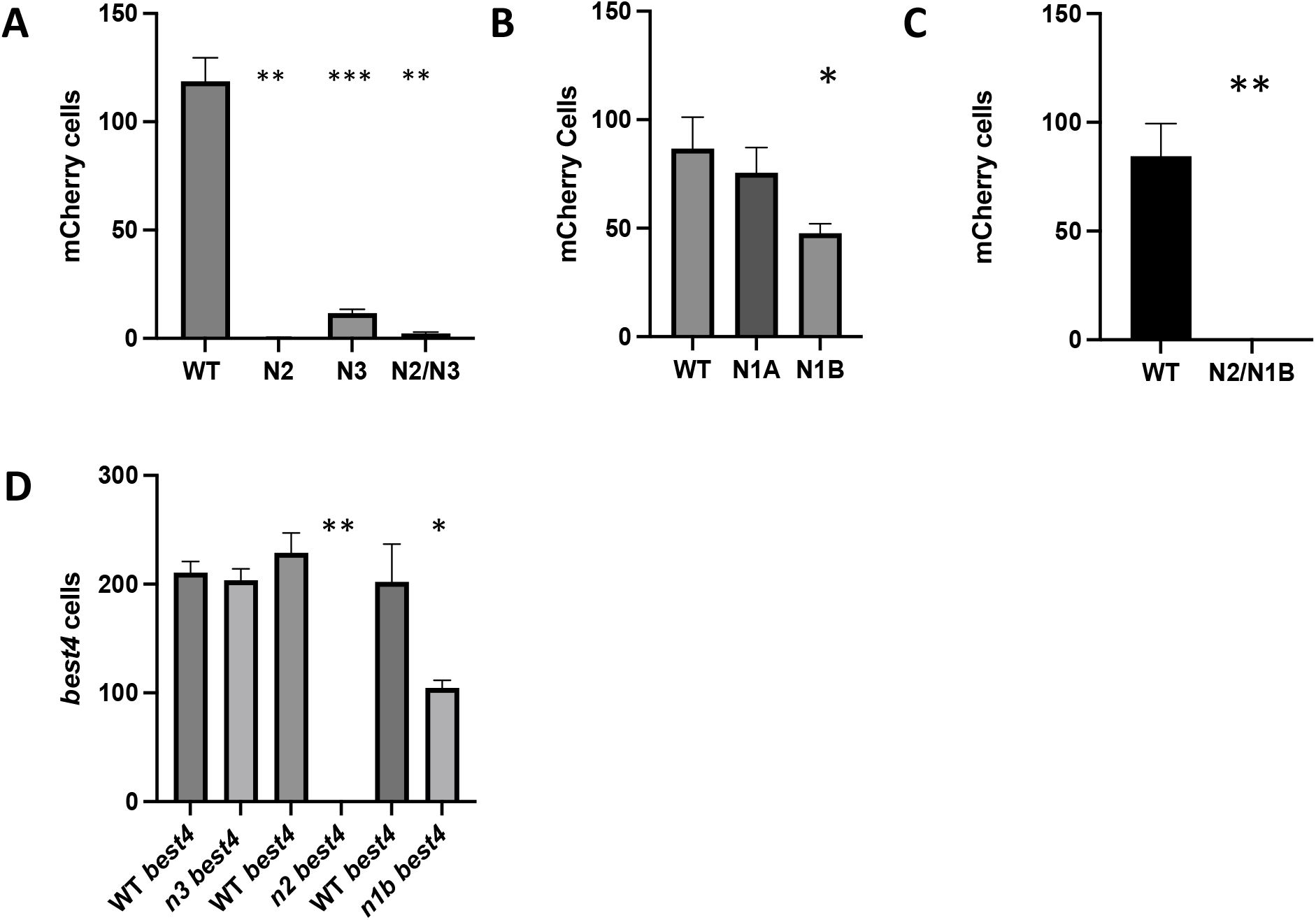
Changes in NRSCs/*best4* cells in mutant Notch receptor mutants. A. Number of nuclear mCherry cells in *notch2, notch3*, and *notch2/notch3* double mutant compared to WT siblings. WT and mutant siblings are paired. B. Number of nuclear mCherry cells in *notch1a* and *notch1b* mutants compared to WT siblings. C. Number of nuclear mCherry cells in notch2/notch1b compared to WT siblings. D. Number of *best4* cells in *notch3, notch2*, and *notch1b* compared to WT siblings. mCherry cells *notch 2, notch3*, and *notch2/3* n=7 to 9; mCherry *notch 1a* and *1b* n=5 to 9; mCherry *notch2/1b* n= 8; *best4* cells n=6 to 10 ^*^ p value ≤ 0.05; ^**^ p value ≤ 0.01; ^***^ p value ≤ 0.001

Loss of nuclear mCherry in *notch2, notch3*, and *notch1b* mutants should also result in loss of *best4* expression. *notch2* mutants have a loss of all *best4* expression within the intestinal epithelium (Figure 3D). *notch1b* mutants have a loss of 54% of *best4* cells compared to WT numbers (Figure 3D). However, *notch3* mutants retain WT levels of *best4* expression (Figure 3D). Loss of all *best4* cells in *notch2* mutants suggests that this receptor is critical for development of the cells. Loss *best4* cells with in both *notch2* and *notch1b* mutants suggests each receptor may be acting on different subtypes.

### Loss of Notch2 results in increased epithelial proliferation

As *notch2* mutants lose all *best4* cells, this should provide an opportunity to identify roles for *best4* cells within the intestinal epithelium. Previously, we identified a role for *best4* cells in epithelial proliferation (Li et al., 2019). We find that interruption of Notch signaling between 74 to 76 hpf (using DAPT) results in increased epithelial proliferation within the anterior intestine but not the posterior when recorded at 5 dpf (Li et al., 2019). The period between 74 to 76 hpf was targeted due to a large number of embryonic *best4* cells differentiating during this period (Li et al., 2019). The only cells with activated Notch signaling during this period appear to be *best4* cells. Disruption of Notch signaling between 74 to 76 hpf should only interrupt development of *best4* cells. With the loss of all intestinal epithelial *best4* cells in embryos with the *notch2* receptor mutation, these mutants should also have increased epithelial proliferation.

To determine whether loss of Notch2 results in increased epithelial proliferation, *notch2* mutants and WT siblings were labeled with EdU at 4 dpf and analyzed at 5 dpf similar to investigations interrupting all Notch signaling using DAPT. As with previous experiments, we analyzed proliferation in an anterior intestinal region as well as a region in the posterior intestine (analyzed regions shown in Figure 1A). While interruption of Notch signaling using DAPT results in increased epithelial proliferation only in the anterior region (Li et al., 2019), we find statistically significant increases in epithelial proliferation in both the anterior and posterior intestine in *notch2* mutants (Figure 4). Increased epithelial proliferation in both anterior and posterior regions of *notch2* mutants may be due to loss of Notch signaling (as observed by lack of reporter) with interruption of development of BEST4+ cells.

**Figure 4.**
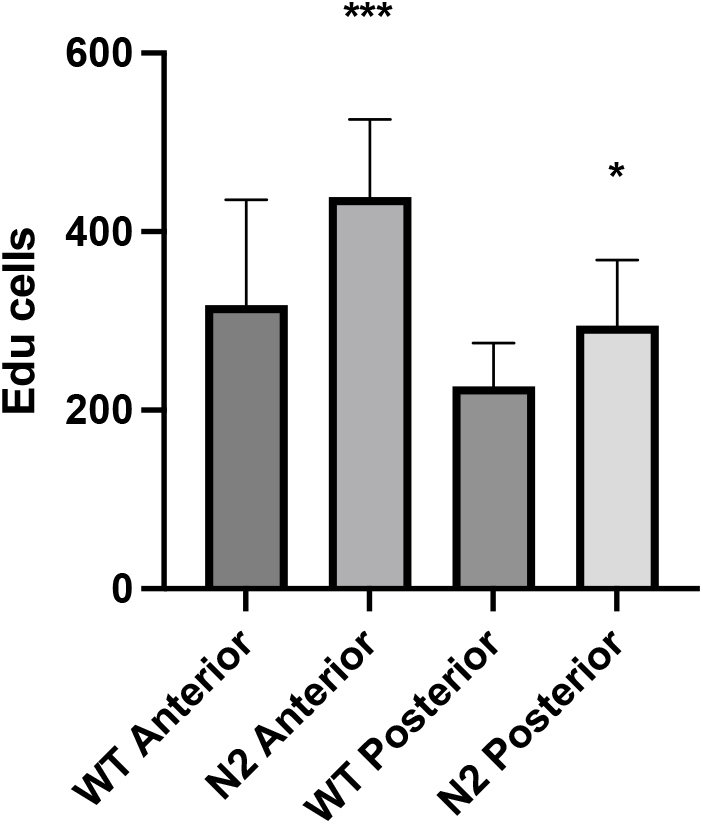
Changes in epithelial proliferation in *notch2* mutants. Change in number of proliferating intestinal epithelial cells comparing *notch2* mutants with WT. WT n=8; *notch2* n=10; ^*^ p value ≤ 0.05; ^***^ p value ≤ 0.001

## Discussion

While it has been shown that Notch signaling is used multiple times during development within the embryonic intestine (Roach et al., 2013), the specific receptors involved in each signaling event have not previously been identified. Here we investigate which receptors are involved in two Notch signaling events during development of the intestine; the choice between secretory and enterocyte cells and development of BEST4+ cells.

The choice between secretory and enterocyte cells begins as future secretory cells upregulate Notch ligands followed by lateral inhibition. Upregulated Notch ligands activate Notch receptors in surrounding cells, inducing the cells to become enterocytes (Capdevila et al., 2021). Disruption of Notch signaling during lateral inhibition results in increased numbers of secretory cells (Roach et al., 2013). During zebrafish embryogenesis, Notch inhibition between 64 to 74 hpf increases the number of secretory cells, suggesting this is the period for the choice between secretory and enterocytes within the embryonic intestinal epithelium (Roach et al., 2013). Using individual Notch receptor mutants, we observe no changes in the number of secretory cells. In contrast, when *notch2* and *notch3* or *notch2* and *notch1b* mutants are combined, we find significant increases in secretory cell numbers, suggesting redundant use of the receptors in the choice between secretory and enterocyte cells.

Redundancy among Notch receptors is also observed in the mammalian intestinal epithelium (Riccio et al., 2008). Loss of individual Notch 1 and Notch 2 receptors produces no change in phenotype. However, loss of both Notch 1 and Notch 2 in the mammalian epithelium results in conversion of the crypt to post mitotic goblet cells (Riccio et al., 2008). While zebrafish and mammals use a different combination of receptors for the choice between secretory and enterocyte cells within the intestinal epithelium, they do share a common theme of redundant action. Redundancy of Notch receptors in the choice between secretory and enterocyte cells allows the same numbers of enterocytes to differentiate with loss of a single Notch receptor, however, there may be alterations in the type of enterocytes that form. Differentiation of particular types of enterocytes may require specific combinations of Notch receptors to create correct ratios of cells. While we identify Notch receptors involved in the choice between secretory and enterocytes, whether there are changes in types of enterocytes will need to be identified by other means.

Notch signaling is used again in development of a group of cells that we previously called Notch Receiving Secretory Cells (NRSCs), which we identify as BEST4+ cells. BEST4+ cells begin developing at 74 hpf following the choice between secretory and enterocyte cells. We previously identified NRSCs with a Notch activated (using the TP1 promoter) CreERT2 transgene to activate a fluorescent transgenic reporter following 4OHT addition(Li et al., 2019). We find that all cells with activated Notch receptors express *best4*. Notch activation in *best4* cells suggests that Notch signaling plays a role in development of these cells. However, we find only about 40% of *best4* cells express the Notch activated fluorescent reporter. BEST4+ cells have differences in regional expression, (Malonga et al., 2024) which is likely to represent different subtypes of the cell population throughout the intestine. We suggest that cells with and without expression of the activated TP1 promoter and fluorescent reporter are two different subtypes of *best4* cells. As described below, we hypothesize that one difference between these potential *best4* cell subtypes is activation of the Notch 3 receptor.

Unlike the choice between secretory and enterocytes, which has redundant use of Notch receptors, we find significant changes in *best4* cells expressing and not expressing the Notch activated reporter following loss of either Notch2 or Notch1b receptors. In *notch 2* mutants there is a loss of almost all cells expressing fluorescent Notch activated reporter and all *best4* cells. Activation of Notch 2 receptor appears to be critical for *best4* cell differentiation. In *notch1b* mutants, 54% of the cells expressing the fluorescent Notch reporter are present and 44% of *best4* cells are present.

While there is loss of *best4* cells both expressing and not expressing the activated Notch fluorescent reporter in *notch1b* mutants, the ratio of these two types of *best4* cells remains similar to WT ratios. In *notch1b* mutants, 45.5% of *best4* cells express the activated Notch fluorescent reporter compared to 36.8 % in WT. Similar ratios of *best4* cell types between *notch1b* mutants and WT, suggest that Notch1b receptor is required for development of a subset of both groups of *best4* cells (expressing and not expressing activated Notch fluorescent reporter). Reduction in both *best4* cell groups in *notch1b* mutants, suggests two additional subgroups of each *best4* cell type: *best4* cells expressing the activated Notch fluorescent reporter that do (1) and do not (2) require Notch1b for development as well as *best4* cells without activated Notch fluorescent reporter that do (3) and do not (4) require Notch1b for development.

Results from loss of *notch3* receptor may explain why more than half of *best4* cells do not express the activated Notch fluorescent reporter. Loss of *notch3* receptor results in loss of most cells expressing the activated Notch fluorescent reporter (reduced to 9.3% of WT) but there is no statistically significant decrease in the number of *best4* cells. With loss of most of the cells expressing the activated Notch fluorescent reporter in *notch3* mutants, we would also expect loss of most of the *best4* cells, similar to losses in *notch2* and *notch1b* mutants.

To explain loss of activated fluorescent Notch reporter expression but not loss of *best4* cells in *notch3* mutants, we suggest that subtypes of *best4* cells have different combinations of activated Notch receptors leading to variable concentrations of Notch Intracellular Domain (NICD). In this scenario, all *best4* cells activate Notch2 receptor but there is another group of *best4* cells that activate both Notch2 and Notch 3 receptors. *best4* cells with only activated Notch 2 receptor would not be able to activate the Notch fluorescent reporter. Cells activating both Notch2 and Notch3 receptors would be the group of *best4* cells that activate the Notch fluorescent reporter. *best4* cells with only activated Notch2 receptor would have a low baseline concentration of NICD while *best4* cells with both activated Notch3 and Notch2 receptors would produce higher concentrations of NICD. *best4* cells with only activated Notch2 receptor would not be able to activate the TP1 promoter, however, these cells would be able to activate low NICD concentration Notch responsive promoters initiating development of the cell type.

Low and high concentrations of NICD between Notch2 only and Notch2/Notch3 *best4* cells could lead to differentiation of different subtypes of *best4* cells. Notch2 only *best4* cells would activate Notch responsive promoters requiring low NICD concentration while Notch2/3 *best4* cells would activate Notch responsive promoters that require both low and high NICD concentrations leading to different *best4* subtypes. Notch3 mutants would then have the same number of *best4* cells, however, all the *best4* cells would be of the subtype that only activates Notch2 receptor and lack expression of the activated Notch fluorescent reporter. It would be likely that the increased numbers of Notch 2 only *best4* cells have an even division of requirements for use of Notch1b signaling.

Previously, we identified increased epithelial proliferation in the anterior intestine following inhibition of Notch signaling by DAPT which inhibits the γ-secretase complex that cleaves the intracellular domain of the receptor (Li et al., 2019). After identifying Notch Receiving Cells as *best4* cells, we determined that inhibition of Notch signaling through DAPT treatment reduces the number of *best4* cells. This suggests that Notch inhibition prevents development of *best4* cells which have a role in regulation of epithelial proliferation. If *best4* cells play a role in epithelial proliferation, *notch2* mutants which have a lack of Notch activation as reported by the TP1 fluorescent reporter combination as well as a total lack of *best4* cells as reported by RNA *in situ* hybridization should also have increased epithelial proliferation. In contrast to only increased proliferation within the anterior intestine of DAPT inhibited embryos (Li et al., 2019), in *notch2* mutants we find increases in epithelial proliferation in both the anterior and posterior intestine. Increased epithelial proliferation throughout the intestine would be expected in *notch2* mutants as there is total loss of *best4* cells. DAPT treatment, on the other hand reduces the number of *best4* cells but seems to leave a number of the cells intact providing opportunity to maintain some level of proliferation regulation. The increased levels of epithelial proliferation in *notch2* mutants is likely to be more representative of the effects of *best4* cell loss.

There has been question as to the lineage origin of BEST4+ cells as to whether they arise from the enterocyte or the secretory cell lineage (Malonga et al., 2024). As mentioned above, in single cell sequencing experiments, Notch2 is expressed within BEST4+ cells (Sur et al., 2023b; Willms et al., 2022). In the lateral inhibition choice between secretory and enterocyte lineages, secretory cells upregulate Notch ligands and activate Notch receptors in surrounding epithelial cells to enter the enterocyte pathway. As a result, cells with high Notch expression might be expected to be of enterocyte origin. With BEST4+ cells, we find that Notch expression observed in these cells is involved in a second signaling event after the choice between secretory and enterocyte cells using lateral inhibition (Li et al., 2019).

BEST4+ cells appear to first enter the secretory pathway lineage and then begin differentiating with a second activation of the Notch2, Notch1b, or Notch3 receptors. We have previously identified the Notch Receiving Secretory cells (NRSCs) as arising from a secretory lineage by colocalization with the pan secretory cell marker AnnexinA4 (2F11 antibody) (Li et al., 2019). The NRSC reporter however is only expressed in 40% of the *best4* cells. To demonstrate that the remaining *best4* cells also arise from the secretory lineage, we colocalize *best4* cells with AnnexinA4 expression. In addition, mutants in *ascl1a* have no secretory cells within the intestinal epithelium (Roach et al., 2013). If *best4* cells are secretory in origin, there should be none of these cells in *ascl1a* mutants. We do find an absence of BEST4+ cells in the intestine of *ascl1a* mutants, further suggesting a secretory origin for the BEST4+ cells.

We find that Notch receptor signaling events within the intestinal epithelium vary between the use of single receptors in development of the *best4* cells and redundant use of Notch receptors with the decision between secretory and enterocyte cells. During embryogenesis, the decision between secretory and enterocyte cells appears to occur during a distinct period followed by differentiation into respective cell types. Initially, BEST4+ cells appear to utilize one or two Notch receptors during differentiation but there may be a more complex interplay between Notch receptors. Activation of different Notch receptors may then result in differentiation of different groups of BEST4+ cells; some with low Notch activation and some with high Notch activation driven by Notch3 activation. We also suggest a novel role for BEST4+ cells in regulation of epithelial proliferation.

## Materials and Methods

### Fish stocks

Fish maintenance and matings were performed as previously described (Westerfield, 1993). Notch mutant fish were genotyped with PCR followed by restriction digestion. Derived Cleaved Polymorphic Sequences (dCAPS) was used to distinguish *notch1b*^*sa11236*^ *(Gerri et al*., *2018)* and *notch3*^*fh332*^ (Quillien et al., 2014) mutants. Forward dCAPS primers for *notch1b*^*sa11236*^ were designed to cut either WT (5’-CCGACTACCTGGGCGGATACACATG-3’; restriction enzyme-PciI) or mutant (5’-CCGACTACCTGGGCGGATACTCATG-3’; restriction enzyme-BsphI). *notch2*^*eI515*^ mutants have a deletion which removes a Cla restriction site(Barske et al., 2016). The *notch1a*^*b420*^ mutant creates an AlfI restriction site (Gray et al., 2001). The *ascl1a* ^*t25215*^ *(Pogoda et al*., *2006)* mutant generates a new Hind III restriction site. Both *notch1a*^*b420*^ and *ascl1a* ^*t25215*^ have primers designed 5’ and 3’ to the difference in restriction sites using NCBI Primer-Blast and Dotmatic’s Snapgene.

### Detection of Notch Receiving Secretory Cells (NRSCs)

To detect Notch receiving secretory cells (NRSCs), the transgene Tg(T2KTp1glob:creERT2)jh12 (referred to as NCre driver) was used to drive expression of inducible CreERT2 in cells that receive Notch signaling (Wang et al., 2011). The transgene drives CreERT2 from 12 RBP-Jκ-binding sites and a minimal β-globin promoter. The Ncre driver is combined with the transgene Tg(T2Kβactin:loxP-stop-loxP-hmgb1-mCherry)jh15 ( referred to as Cre responder) to mark NRSCs with nuclear mCherry expression (Wang et al., 2011). Induction of CreERT2 with 4-Hydroxytamoxifen (4OHT, T176, Sigma) for a 2-h period was performed at the same concentration (5 μM) as previously reported, diluted from a stock concentration of 10mM dissolved in 100% ethanol (Wang et al., 2011). Following induction, NRSCs and their progeny continuously express nuclear mCherry.

### HCR and immunohistochemistry

*best4* RNA *in situ* hybridization was performed using a probe synthesized by Molecular Instruments and detected with HCR RNA-FISH kit following the standard protocol. Immunohistochemistry with AnnexinA4 (2F11 Abcam ab71286) and mCherry (Takara DsRed polyclonal antibody 632496) was incubated overnight followed by 45 minutes of PBST washes, incubation of secondary antibody (Invitrogen Alexa Fluor 488 and 594) for 2 hours, followed by 45 minutes wash PBST. For both procedures, 5 dpf embryos were permeabilized with 15.5 ug/ml Proteinase K (Sigma, P2308) after rehydration in PBST.

### Imaging

Intestines imaged in whole mount were dissected using #5 Dumont Tweezers from the specimen and intestines were mounted on a slide in Vectashield (Vector Laboratories). Specimens were then imaged with a Leica TCS SP8 X confocal.

### EdU incorporation and detection

Embryos and larvae are injected with a microcapillary needle in the intestinal region (either directly into the intestine or peritoneal space) with 2.5mM 5-ethynyl-20-deoxyuridine (EdU). EdU injected individuals were injected at 4 dpf and grown to 5 dpf. Embryos were fixed in 4% formaldehyde, 2 h to overnight. Incorporation of EdU was detected following the standard protocol for the 488 Click-it Kit imaging kit (Invitrogen).

### Statistical Analysis

Determination of significant differences between control and experimental samples was identified using Student’s t-test. Results with P values of ≤0.05 were considered statistically significant. Statistically significant values are indicated by asterisks in the bar graphs. The number of asterisks indicate increasing significant differences with lower P values indicated in the respective legend.

## Acknowledgments

The authors would like to thank Gage Crump for providing the *notch2* mutation, Cecilia Moens for the *notch 3* mutant, and Christine Beattie for the *notch1a* mutant. We thank Michael Parsons for providing *Tg(T2KTp1glob:creERT2)jh12* and *Tg(T2Kβactin:loxP-stop-loxPhmgb1-mCherry)jh1, Tg[nkx2*.*2a:mEGFP]*. Research reported in this publication was supported by the Eunice Kennedy Shriver National Institute of Child Health & Development of the National Institutes of Health under Award Number 1R15HD108689-01. The content is solely the responsibility of the authors and does not necessarily represent the official views of the National Institutes of Health.

